# Natural Variation Meets Synthetic Biology: Promiscuous Trichome Expressed Acyltransferases from *Nicotiana acuminata*

**DOI:** 10.1101/2022.02.25.482030

**Authors:** Craig A. Schenck, Thilani M. Anthony, MacKenzie Jacobs, A. Daniel Jones, Robert L. Last

## Abstract

Acylsugars are defensive, trichome-synthesized sugar esters produced in plants across the Solanaceae (nightshade) family. Although assembled from simple metabolites and synthesized by a relatively short core biosynthetic pathway, tremendous within- and across-species acylsugar structural variation is documented across the family. To advance our understanding of the diversity and the synthesis of acylsugars within the *Nicotiana* genus, trichome extracts were profiled across the genus coupled with transcriptomics-guided enzyme discovery and *in vivo* and *in vitro* analysis. Differences in the types of sugar cores, numbers of acylations, and acyl chain structures contributed to over 300 unique annotated acylsugars throughout *Nicotiana*. Placement of acyl chain length into a phylogenetic context revealed that an unsaturated acyl chain type was detected in a few closely-related species. A comparative transcriptomics approach identified trichome-enriched *Nicotiana acuminata* acylsugar biosynthetic candidate enzymes. > 25 acylsugar variants could be produced in a single enzyme assay with four acylsugar acyltransferases (NacASAT1-4) together with structurally diverse acyl-CoAs and sucrose. Liquid chromatography coupled with mass spectrometry screening of *in vitro* products revealed the ability of these enzymes to make acylsugars not present in *Nicotiana* plant extracts. *In vitro* acylsugar production also provided insights into acyltransferase acyl donor promiscuity and acyl acceptor specificity as well as regiospecificity of some ASATs. This study suggests that promiscuous *Nicotiana* acyltransferases can be used as synthetic biology tools to produce novel and potentially useful metabolites.

**ONE SENTENCE SUMMARY:** Analysis of *Nicotiana* glandular trichome metabolites and BAHD acyltransferases revealed diverse sucrose and glucose based acylesters.

## INTRODUCTION

Plants produce a vast array of small molecules, many of which are classified as specialized metabolites (Wink, 2010; Jacobowitz and Weng, 2020). Despite being assembled from simple building blocks, specialized metabolites can be structurally diverse due to lineage-specific biosynthetic enzymes (Fang et al., 2019). Unlike the ubiquitous products of core metabolism such as nucleotides and amino acids, specialized metabolites display restricted taxonomic distribution and are confined to certain cell or tissue types (Pichersky and Lewinsohn, 2011; Beaudoin and Facchini, 2014; Schenck and Last, 2020). While the function(s) of most specialized metabolites remain unknown, there are an increasing number of examples of metabolites that enable both positive (e.g., pollinator attraction) and negative (e.g., herbivore deterrence) plant-environmental interactions (Howe and Jander, 2008; Massalha et al., 2017; Weng et al., 2021). Additionally, specialized metabolites have been adopted and modified by humans to serve medicinal, nutritional, and industrial roles (Pichersky et al., 2006; Schramek et al., 2010; Srinivasan, 2016; Weng et al., 2021). Plants have evolved various anatomical structures for specialized metabolite synthesis and storage, which serve to prevent damage and competition with other metabolic pathways (e.g. laticifers Schilmiller et al., 2012b; Schuurink and Tissier, 2020). Glandular trichomes are a common example of a biological factory for producing various classes of metabolites including terpenoids (Sallaud et al., 2009; Pichersky and Raguso, 2018), flavonoids (Schmidt et al., 2011; Kim et al., 2014) and acylsugars (Schilmiller et al., 2010; Fan et al., 2019; Schuurink and Tissier, 2020).

Sugar esters, known as acylsugars, represent a class of specialized metabolites widely distributed across the Solanaceae (nightshade) family (Balcke et al., 2017; Chortyk et al., 1997; Fan et al., 2019; King and Calhoun, 1988; Kroumova et al., 2016; Luu et al., 2017; Moghe et al., 2017; Nakashima et al., 2016), and in other families, including Caryophyllaceae and Convolvulaceae (Pereda-Miranda et al., 2010; Kruse et al., 2021). Within the Solanaceae, acylsugars accumulate in glandular trichomes and are involved in defense against an array of biological pests (Leckie et al., 2016; Ghosh and Jones, 2017; Luu et al., 2017; Smeda et al., 2017; Feng et al., 2021; Kortbeek et al., 2021). Sugars and acyl-CoAs from core metabolism are acylsugar precursors, with the acyl-CoAs derived from branched chain amino acid (BCAA) or fatty acid metabolism (Kandra et al., 1990; Slocombe et al., 2008; Chang et al., 2020; Fan et al., 2020; Mandal et al., 2020). While acylsugars with sucrose cores are common across the Solanaceae, other sugar core types are found. Acylglucoses accumulate in some genera including *Solanum* and *Nicotiana* (Matsuzaki et al., 1989; Leong et al., 2019; Lou et al., 2021) and inositol-based esters have also been detected in the *Solanum* (Herrera-Salgado et al., 2005; Leong et al., 2020; Lou et al., 2021).

Acyl chain lengths, branching patterns and composition are additional contributors to acylsugar structural variation and are determined, at least in part, by the acyl-CoA pool in the trichomes (Kandra et al., 1990; Slocombe et al., 2008; Fan et al., 2020). Acyl-CoAs longer than four and five carbons can be elongated, leading to varied building blocks for acylsugar biosynthesis (Walters and Steffens, 1990; Kroumova and Wagner, 2003). Structurally varied acyl-CoA and sugar precursors contribute to a vast number of annotated acylsugar types found throughout the Solanaceae (Moghe et al., 2017; Fan et al., 2019) and in combination with various positions and numbers of acylations provide the potential for production of an almost unlimited number of acylsugar structural variants.

The core trichome acylsugar pathway is catalyzed by a series of clade III BAHD acyltransferase family enzymes (D’Auria, 2006), which esterify acyl groups from acyl-CoAs onto sugar core oxygen atoms (Schilmiller et al., 2012a; Schilmiller et al., 2015; Fan et al., 2016). In the cultivated tomato *S. lycopersicum* M82, trichome-enriched acylsugar acyltransferases (ASATs) sequentially add four acyl chains at specific positions on a sucrose core; these are named SlASAT1-SlASAT4 reflecting their order in the pathway (Schilmiller et al., 2012a; Schilmiller et al., 2015; Fan et al., 2016). Comparative transcriptomics coupled with phylogenetics, *in vitro* biochemistry and *in planta* gene silencing studies identified homologous ASATs from species across the Solanaceae with functions reminiscent of those from tomato (Moghe et al., 2017; Nadakuduti et al., 2017; Leong et al., 2020; Lou et al., 2021). However, differences in acylation order (Fan et al., 2017) and ASAT gene loss/gain events (Moghe et al., 2017) contribute to distinct biosynthetic pathways leading to acylsugar structural variation across the Solanaceae.

The complex mixtures of acylsugars produced across the *Nicotiana* genus have led to its use as an experimental model to test the roles of these metabolites in adaptation, including in direct and indirect defense (Buta et al., 1993; Puterka et al., 2003; Weinhold and Baldwin, 2011; Luu et al., 2017; Feng et al., 2021). Reduction of acylsugars through transgenic approaches targeting acylsugar biosynthetic genes demonstrate the roles of acylsugars in protection against insects, fungi, and desiccation (Luu et al., 2017; Feng et al., 2021). *Nicotiana* acylsugars with long acyl chains had greatest antibiotic activity (Chortyk et al., 1993) and eight carbon long acyl chains were most protective against white whiteflies (Chortyk et al., 1996), demonstrating that acylsugar structural variants have differing biological activities. Acylsugars also have industrial applications (Garti et al., 2000), and acyl chain types contribute to this activity. Exploration of acylsugar structural diversity across *Nicotiana* species and identification of biosynthetic enzymes with diverse activities provides the opportunity to produce specific acylsugar types with enhanced biological and industrial activities.

We describe analytical chemical screening of *Nicotiana* trichome metabolites that revealed hundreds of acylsugar structures that differ in sugar core composition, acylation position and numbers, branching pattern and lengths of acyl chains. Correlation of metabolite data with phylogenetic distribution revealed lineage-specific acylsugars. Sugar acyltransferase enzymes identified using a comparative transcriptomics approach were characterized by *in vitro* biochemistry. *N. acuminata* acyltransferases produced a wide variety of products, including those not found in *Nicotiana* acylsugar extracts, highlighting the metabolic promiscuity of these enzymes. *In vitro* acylsugar combinatorial analysis with *Nicotiana* acyltransferases provides a step towards a synthetic biology platform aimed at production of acylsugars with varied biological activities.

## RESULTS

### Acylsugar diversity within the *Nicotiana* genus

To analyze trichome acylsugar structural variation across the *Nicotiana* genus, surface metabolites were extracted from leaves of 6-week-old plants and screened using complementary analytical approaches (Figure 1). Reverse phase liquid chromatography (RP-LC) coupled with quadrupole time-of-flight mass spectrometry (QTOF-MS) revealed total numbers of chromatographically separated and detectable acylsugars per species and provided information about numbers and lengths of acyl chains on individual acylsugars. LC-MS with collision-induced dissociation (CID) resolved the type and abundance of sugar cores; GC-MS provided information about acyl chain structure and abundance; and nuclear magnetic resonance spectroscopy (NMR) was used to resolve the absolute structures of two *Nicotiana* acylsugars to establish acylation positions.

**Figure 1.**
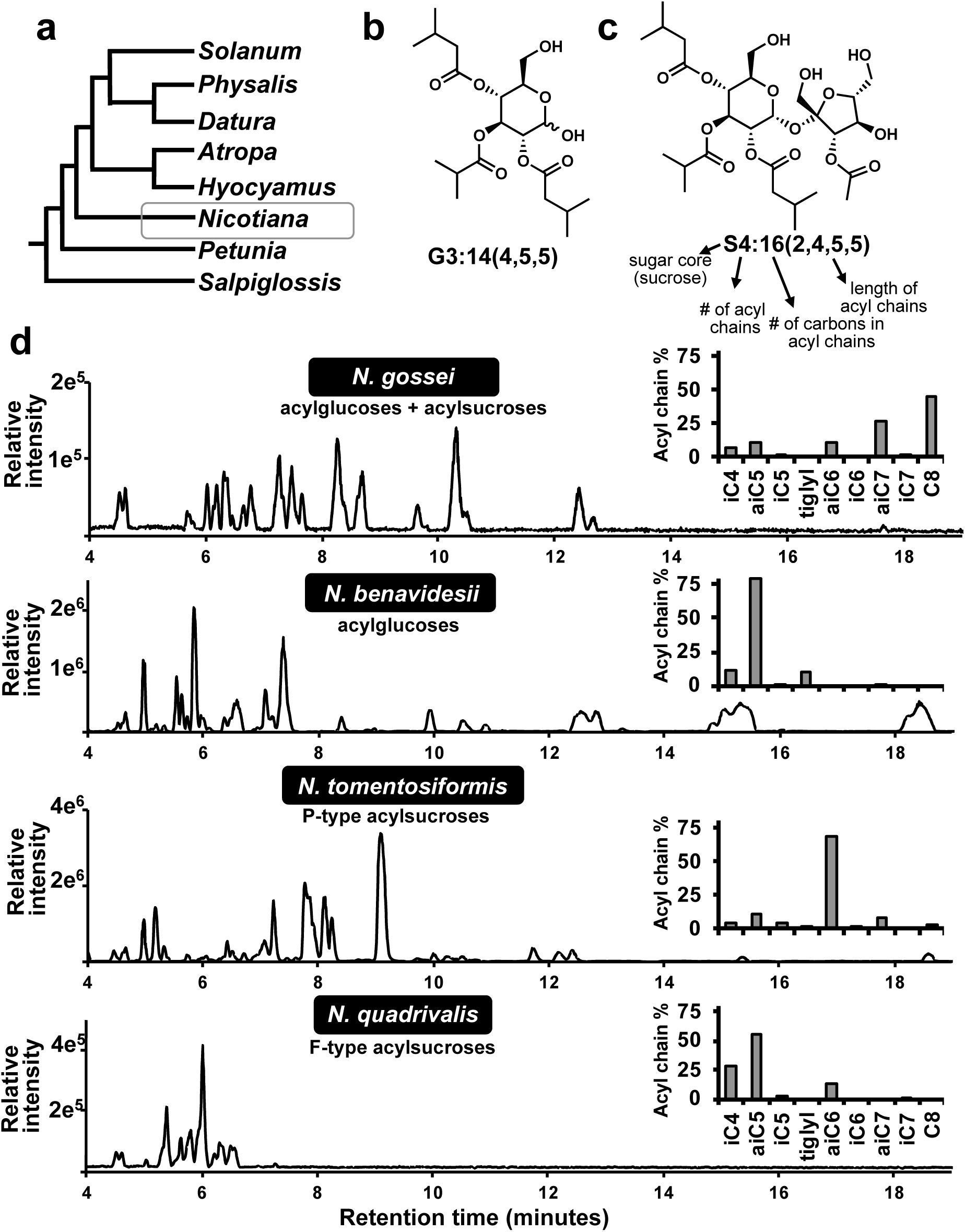
Acylsugar structural diversity in *Nicotiana.* (a) Acylsugar screening in the *Nicotiana* genus (highlighted in gray) within the Solanaceae family. Examples of acylglucose **(b)** and acylsucrose **(c)** structures found in multiple *Nicotiana* species. Acylsugar extracts were analyzed using LC-MS **(d)** and acyl chain structural variation was determined through GC-MS (insets in d). Acylsugars were chromatographically separated and analyzed in negative ion mode **(d)**. These four species represent examples of the structural variation of *Nicotiana* acylsugar extracts. Specifically, *N. benavidesii* contains only acylglucoses, *N. gossei* contains a mix of acylsucroses and glucoses. *N. tomentosiformis* and *N. quadrivalis* contain only acylsucroses but differ in the location of the acetylation. *N. tomentosiformis* has the acetylation on the glucose ring (P-type) whereas *N. quadrivalis* has the acetylation on the fructose ring (F-type).

### LC-MS annotation of acylsugar structural variants

LC QTOF-MS was used to annotate acylsugars based on accurate mass measurements of unfragmented ions and fragmentation patterns, providing evidence for the numbers of detectable acylsugars in each species. Collision-induced dissociation (CID) LC-MS QTOF in negative-ion mode yielded information about the lengths and numbers of acyl chains on individual sugar cores (Figure 1 & Supplemental Figure S1; Ghosh et al., 2014). The complementary approach of positive-ion mode CID was used to infer acyl chain position on the pyranose (six-membered) or furanose (five-membered) ring of acylsugars with a disaccharide core (Supplemental Figures S2 & S3), as the cleavage of the glycosidic linkage without ester cleavage yielded diagnostic fragment ion masses. *N. acuminata* extracts contained 21 annotated acylsugars, while *N. forgetiana* contained 19 annotated acylsugars plus five with abundances too low to be confidently annotated based on fragmentation pattern (Supplemental Table S1). In contrast, some acylsugar-producing *Nicotiana* species accumulated far fewer abundant acylsugars: for example, *N. knightiana* accumulated only four annotatable acylsugars (Supplemental Table S1). Finally, some of the *Nicotiana* species screened did not produce detectable acylsugars, which correlated with lack of visible leaf and stem glandular trichomes (Supplemental Figure S4). Acylsugars were observed with as few as two, and as many as five acylations: the difference in numbers of acylations represented a major source of structural variation across the genus (Supplemental Table S1).

Despite the relatively close evolutionary relationships of the species screened, the majority of acylsugars were detected in a single genotype, with only 21 acylsugars detected in more than one species (Supplemental Table S1). For example, an acylsugar with a retention time of 4.63 minutes and 681.29 *m/z* of the formate adduct was detected in both *N. glauca* and *N. occidentalis* (Supplemental Table S1) and annotated to have a disaccharide core (determined to be sucrose through base hydrolysis; Supplemental Figure S5). This tetraacylsugar had chains of 2, 5, 5, and 5 carbons long (Supplemental Table S1), with the acetylation on the furanose ring. An acylsugar with such a composition is referred to as S4:17(2^F^,5,5,5) where S indicates sugar core type (sucrose in this case) with 4 acylations: totaling 17 carbons and the individual acyl chain lengths of 2, 5, 5, and 5 carbons. When additional information can be inferred, such as location of an acetyl group on a pyranose or furanose ring, this is indicated with a superscript P or F. Of these 21 acylsugars, only two were identified in more than two species, both with an annotation of G3:15(5,5,5), but different retention times, suggesting that they are isomers (Table S1). G3:15 isomers were detected in closely related species, which formed a clade in the *Nicotiana* phylogenetic tree (Supplemental Figure S1).

We also documented cases where metabolites with the same accurate mass accumulated in multiple species but appear to be positional isomers. For example, a negative mode molecular ion with *m/z* 653.27 was identified in five different species with a similar fragmentation pattern, but different chromatographic elution times (Supplemental Table S1). Fragmentation patterns revealed that the acylsugars of *m/z* 653.27 are disaccharides with four acylations of carbon lengths 2, 4, 4, and 5. Evidence for the hypothesis that these chromatographically separable tetraacylsugars differ in esterification positions was obtained using positive ion mode. In *N. kawakamii,* S4:15(2^P^,4,4,5) was acetylated on the pyranose ring and a retention time of 4.10 minutes, whereas in *N. obtusifolia* and *N. linearis* S4:15(2^F^,4,4,5) had the acetylation on the furanose ring and a retention time of 4.55 minutes (Supplemental Table S1). Evidence that furanose ring acylated S4:15(2^F^,4,4,5) in *N. quadrivalis* and *N. africana* is a third positional or structural isomer is from its retention time of 5.40 minutes (Supplemental Figure S1). The combination of retention time, ring position acetylation and fragmentation pattern contributed to annotation of greater than 300 acylsugar structural variants across the entire genus with many more of abundance too low for confident annotation.

### Nicotiana sugar core analysis

Differences in sugar core type was another source of acylsugar structural variation. LC-MS QTOF of intact acylsugars provides indication of whether the sugar core is a mono or disaccharide but does not readily distinguish isomeric sugar cores. We characterized the types and quantities of *Nicotiana* acylsugar cores by base hydrolysis of the acyl chains following free sugar removal by phase partition. Analysis of these products using LC with a polar stationary phase and targeted MS/MS revealed diversity in sucrose/glucose ratios (Supplemental Figure S5) including species with only sucrose or glucose cores detected (e.g., *N. langsdorfii* and *N. paniculata,* respectively), and species with mixtures of sucrose and glucose cores (e.g., *N. rotundifolia*; Supplemental Figure S5). Sucrose-based acylsugars were the predominant types detected, consistent with published results from a subset of *Nicotiana* species (Supplemental Figure S5; Arrendale et al., 1990; Kroumova et al., 2016; Luu et al., 2017; Matsuzaki et al., 1989). Placement of acylsugar core type and composition onto a *Nicotiana* phylogenetic tree showed that closely related species tended to have similar cores: for example, sucrose cores were exclusively detected in almost all species centered on *N. glauca* (Supplemental Figure S5). In contrast, acylglucose-producing species were enriched in a single clade (Supplemental Figure S5).

### Phylogenetic distribution of acyl chain structural variation

To gain insight into the phylogenetic distribution of acylsugar chain lengths, acyl chains revealed by LC-MS CID fragmentation patterns were mapped onto a *Nicotiana* phylogeny (Supplemental Figure S1). Notably, no acyl chains longer than eight carbons were detected (Supplemental Figure S1), consistent with previously published acylsugar analysis within *Nicotiana* and closely related species such as *P. axillaris* and *S. sinuata* (Matsuzaki et al., 1989; Kroumova et al., 2016; Moghe et al., 2017; Nadakuduti et al., 2017). This is in contrast to other Solanaceae lineages, which have acyl chain lengths up to 14 carbons; including species within the *Solanum, Hyoscyamus,* and *Physalis* genera (Ghosh et al., 2014; Zhang et al., 2016; Moghe et al., 2017; Fan et al., 2019). C5 acyl chains derived from leucine or isoleucine were detected in all tested species except *N. benthamiana* (Supplemental Figure S1), whereas longer acyl chains were more sporadically distributed (Supplemental Figure S1). Phylogenetic placement of metabolite data revealed that acetylated acylsugars were found across the *Nicotiana*, except for the clade containing *N. paniculata* (Supplemental Figure S1).

We obtained evidence for C5 and C6 acyl chains with a single desaturated C-C bond in a subset of species. For example, a C5 desaturated acyl chain accumulated only in *N. acuminata* and closely related species based on presence of *m/z* 99.05 in negative CID mode and consistent neutral losses. *N. tomentosiformis* was the only species that accumulated a detectable acyl chain with *m/z* 113.06 and neutral losses consistent with a desaturated C6 acyl chain (Supplemental Figure S1). While LC-MS CID fragmentation patterns provided information about the mass of the fragments lost, other analytical methods were needed to elucidate the branching patterns and structures of acyl chains.

### Acyl chain structural variations probed by GC-MS

LC-MS CID fragmentation patterns suggested the presence of C5 and C6 desaturated acyl chains. To determine the abundance and structures of the acyl chains, GC-MS analysis was performed on a subset of species with a focus on those that contained a putatively desaturated C5 acyl chain. Branching pattern isomers, and the structures of desaturated acyl chains were identified from the corresponding ethyl esters following acyl chain cleavage from the sugar core (Figure 2 & Supplemental Figure S6). Subterminally branched anteiso (ai) acyl chains were more abundant compared to terminally branched iso (i) acyl chains in most species screened. In *N. megalosiphon*, aiC5, aiC6, and aiC7 acyl chains were all more abundant than their terminally branched isomer (Supplemental Figure S6), consistent with greater contributions of isoleucine than leucine precursors. Library matching of mass spectra and retention times putatively identified the desaturated C5 acyl chain as a tiglyl-ester, an unsaturated five-carbon CoA ester, which is an intermediate in isoleucine catabolism (Basey and Woolley, 1973; Binder, 2010). While the GC-MS and LC-MS results were generally comparable, minor amounts of tiglyl ethyl esters were detected in *N. benavidesii* (Supplemental Figure S6), in contrast to LC-MS data from the same species (Supplemental Table S1). The relative amounts of tiglyl acyl chains compared to all acyl groups determined through GC-MS analysis ranged from 10 % in *N. corymbosa* to 75 % in *N. acuminata* (Figure 2 & Supplemental Figure S6).

**Figure 2.**
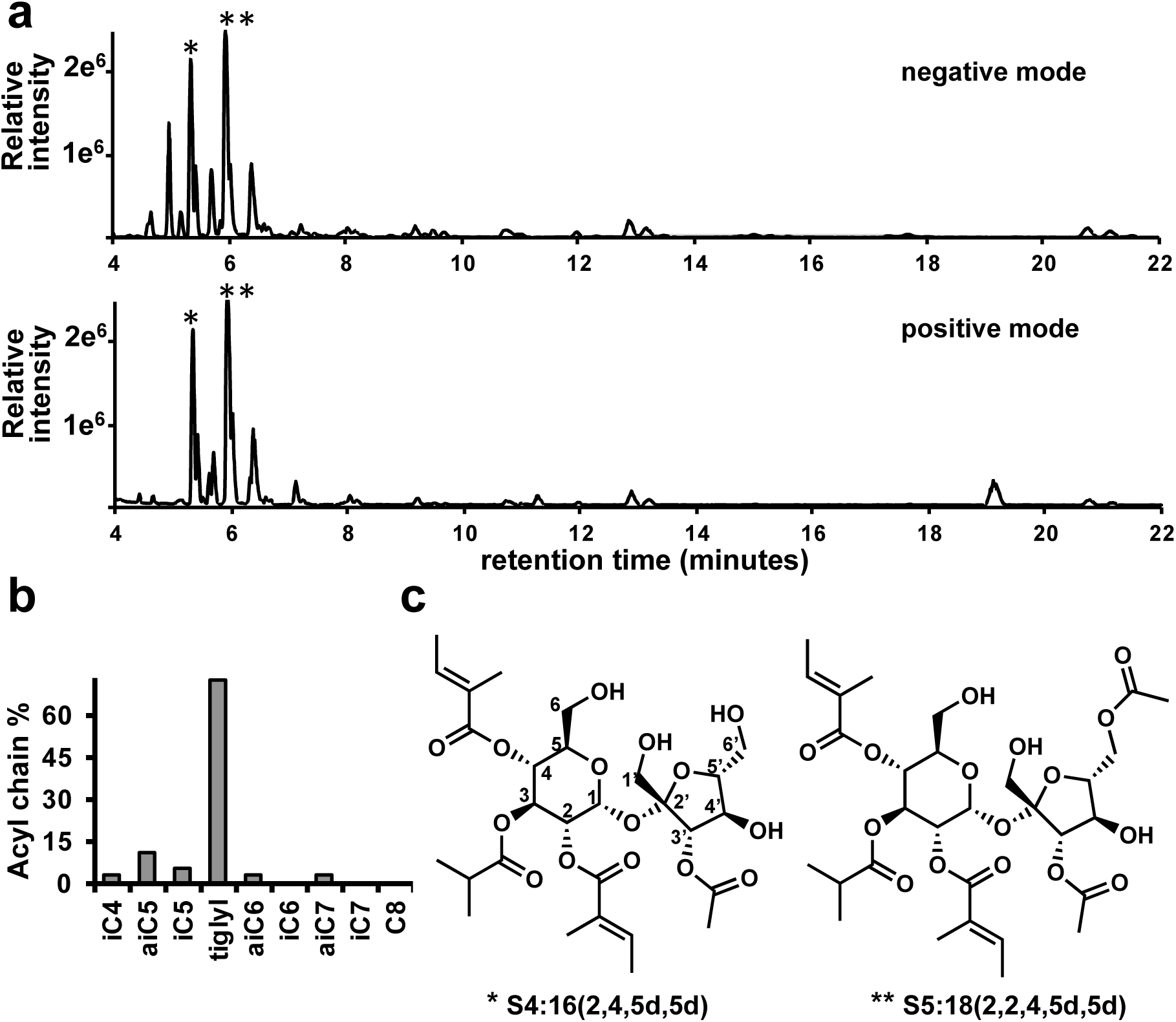
Tiglylated acylsugars in *Nicotiana acuminata.* Acylsugar extracts from *N. acuminata* were analyzed using LC-MS **(a)** and GC-MS **(b)** and structures of two acylsugars (marked by one or two asterisks above the peak, respectively) were resolved using NMR **(c)**. **(a)** Acylsugars were chromatographically separated using a C18-reverse phase column, almost all peaks in the chromatograms correspond to acylsugars. **(b)** Acyl chain types were determined using GC-MS. **(c)** The structures of two tiglyl-chain containing acylsugars were resolved using NMR. NMR analysis using ^1^H, ^13^C, COSY, HSQC, HMBC, TOCSY, and NOESY experiments (Supplemental Data File 1).

### NMR resolved structures of tiglyl acyl chain containing acylsugars

NMR spectroscopy was used to elucidate the structures of acylsugars containing the uncommon tiglyl acyl chain. Two abundant acylsucroses from *N. acuminata* were purified and characterized: one tetraacylated and the other pentaacylated. Each consisted of a sucrose core with tiglyl acylations at the 2- and 4- positions of the pyranose ring (Figure 2c, Supplemental Data File 1), while the pyranose 3- and furanose 3′-positions had iso-branched C4 (iC4) and acetyl groups, respectively (Figure 2c, Supplemental Data File 1). The difference between the two acylsugars was an additional acetyl group at position 6′ of the pentaacylsucrose (Figure 2c, Supplemental Data File 1); the locations of the two acetyl groups and the short, branched acyl chains are reminiscent of the previously resolved acylsugar structures from *S. sinuata* (Moghe et al., 2017).

### Tissue-specific transcriptomics enabled acylsugar acyltransferase discovery

The striking numbers of acylsugars detected across the *Nicotiana* genus offers an opportunity to expand the toolkit of acyltransferase enzyme activities and harness their potential for production of acylsugar structural variants. We focused on *N. acuminata* as a source for candidate enzymes because of its larger number of detected acylsugars including the presence of tiglylated acylsugars. We sought trichome-enriched BAHD acyltransferases, as these likely use different acyl donor substrates compared to other species based on the numbers of acyl chain types detected on *N. acuminata* acylsugars, but did not focus on the enzymes that control acyl-CoA pools.

Trichomes scraped from *N. acuminata* stem and shaved stem tissue were used for RNA-seq-mediated *de novo* transcriptome assembly analysis (Figure 3). To identify ASAT candidates we used an approach successfully employed in species across the family. First, we sought trichome expression enriched homologs of ASATs from tomato, *P. axillaris* and *S. sinuata* (Fan et al., 2016; Moghe et al., 2017; Nadakuduti et al., 2017) by BLAST analysis against the *N. acuminata de novo* assembly. Amino acid sequence alignments with the candidate genes and biochemically characterized ASATs demonstrated the presence of two conserved domains characteristic of functional BAHD acyltransferase enzymes. Four trichome enriched (Figure 3b)

**Figure 3.**
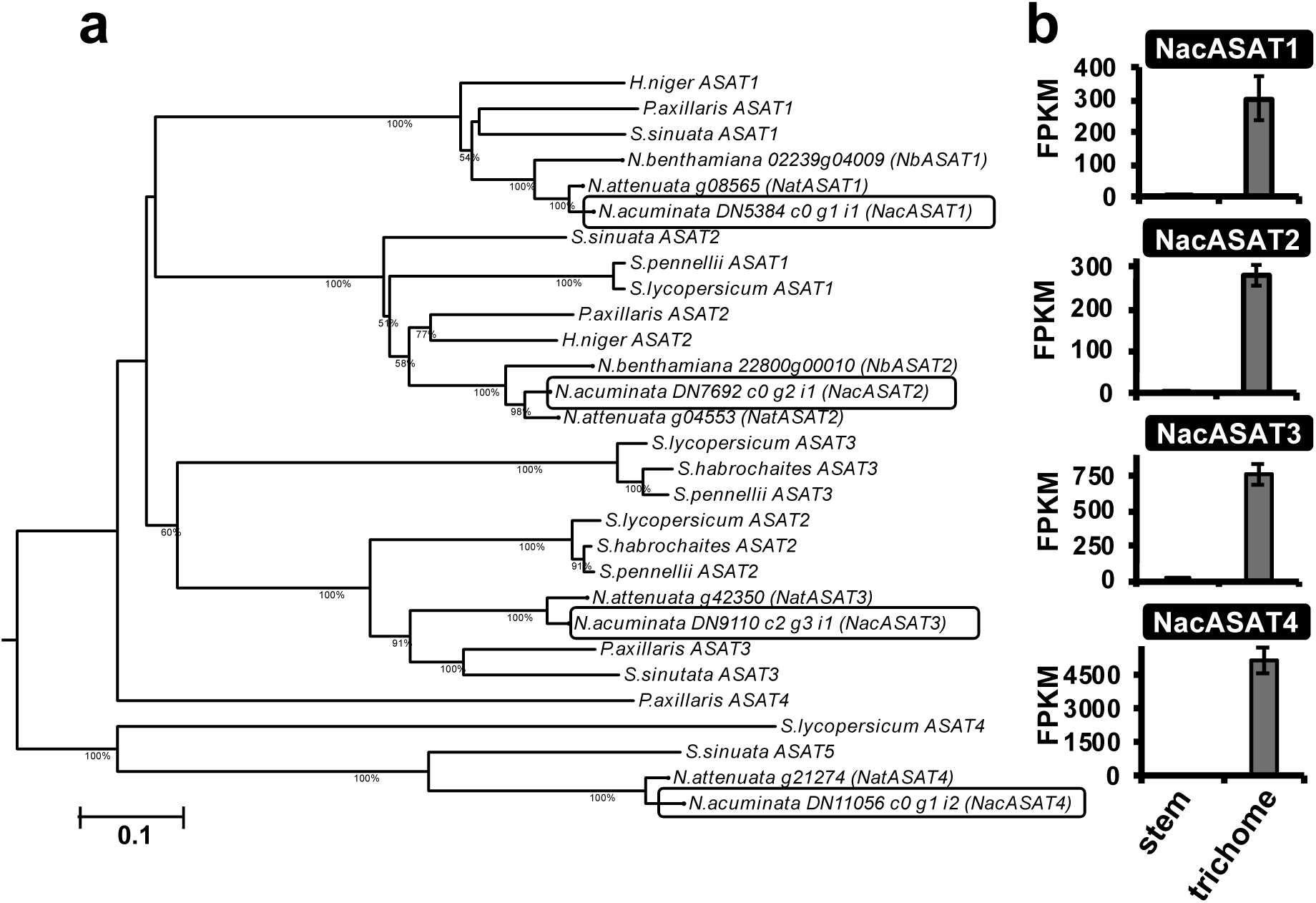
Comparative transcriptomics analysis identifies candidate acyltransferase enzymes. A *de novo* transcriptome assembly was assembled from *N. acuminata* trichome and shaved stem tissue. **(a)** Phylogenetic analysis of biochemically characterized BAHD acyltransferase enzymes involved in acylsugar biosynthesis together with candidate ASATs from *N. acuminata* and some *Nicotiana* species with genome sequence data. ASAT candidates from *N. acuminata* are highlighted in gray boxes. The tree was constructed using maximum likelihood calculated with 1,000 bootstrap values. Bootstrap values less than 50% are not shown. Scale bar represents number of amino acid changes per site. The tree was annotated using iTOL (Letunic and Bork 2019). **(b)** NacASATs are highly expressed in trichome, but not shaved stem tissue. Fragments per kilobase million (FPKM) is shown as the average of three biological replicates ± SEM to represent the gene expression of each candidate ASAT from the RNA-seq dataset.

*N. acuminata* candidates possessed the HxxxD and DFGWG motifs involved in catalysis and substrate binding, respectively (Supplemental Figure S7; D’Auria, 2006). Phylogenetic analysis of the candidate genes showed that each grouped within one of the clades containing previously characterized ASATs (Figure 3a). Candidate ASATs from *N. acuminata* were named based upon phylogenetic grouping with previously characterized ASATs, but do not necessarily reflect their acylation order. *N. acuminata* DN5384_c0_g1_i1 (NacASAT1 ortholog) grouped within a clade containing enzymes from early diverging Solanaceae lineages previously demonstrated to catalyze acylation of non-esterified sucrose using an acyl-CoA (Moghe et al., 2017; Nadakuduti et al., 2017). A NacASAT2 ortholog (DN7692_c0_g2_i1) grouped within a clade that contained ASAT2 enzymes from early diverging Solanaceae lineages, as well as *Solanum* ASAT1s capable of acylating free sugars (Figure 3a, Fan et al., 2016; Moghe et al., 2017). The clade containing NacASAT3 ortholog DN9110_c2_g3_i1 includes biochemically characterized ASAT3s from *P. axillaris* and *S. sinuata* (Moghe et al., 2017; Nadakuduti et al., 2017) and *Solanum* ASAT2s (Fan et al., 2016; Fan et al., 2017). *N. acuminata* DN11056_c0_g1_i2 (NacASAT4 ortholog) grouped within a clade containing acyltransferases previously demonstrated to catalyze acylsugar acetylation (Figure 3a; Fan et al., 2016; Moghe et al., 2017). Taken together, the phylogenetic relationships to previously characterized ASATs, presence of conserved protein domains, and trichome enrichment of *N. acuminata* ASAT homologs led us to characterize their activities.

### In planta and in vitro analysis of N. acuminata ASAT1

A virus induced gene silencing (VIGS) approach was used to test the *in planta* role of the *N. acuminata* ASAT1 ortholog in acylsugar biosynthesis (Supplemental Figure S8a). Two independent VIGS fragments targeting different regions of *NacASAT1* each resulted in a statistically significant decrease in total acylsugar levles compared with an empty vector control (Supplemental Figure S8b). RNA extracted from the same leaves used for acylsugar quantification revealed that *NacASAT1* expression levels were reduced by approximately 35% (Supplemental Figure S8c). These data are consistent with the hypothesis that *NacASAT1* is involved in acylsugar biosynthesis *in planta*.

We conducted *in vitro* biochemical analysis of NacASAT1 to complement the *in planta* approach. A single monoacylated product was observed when the recombinant NacASAT1 was incubated with iC5-CoA acyl donor and sucrose acyl acceptor substrates (Figure 4a). The variety of acyl chains observed in acylsugars found across the *Nicotiana* genus led us to test the ability of NacASAT1 to accept other acyl-CoA substrates ranging from aliphatic iC4 to straight-chain nC14, along with the aromatic donor benzoyl-CoA (Figure 4a). NacASAT1 used all the CoA substrates tested, while preferring longer straight chain CoAs over short-branched CoAs (Figure 4b). An approximately 10-fold difference in activity was observed between the most and least preferred substrates (nC8 and iC4, respectively; Figure 4). We tested acceptor substrate specificity with a range of sugars. As shown in Figure S9, of the four disaccharides tested, NacASAT1 was most active with sucrose, while only minor activity was detected with the glucose dimer trehalose -- approximately 2% of sucrose, which is composed of glucose and fructose. No activity was detected with monosaccharides or *myo*-inositol (Supplemental Figure S9). Taken together, these data reveal broad acyl donor promiscuity, and strong acylsucrose acyl acceptor specificity.

**Figure 4.**
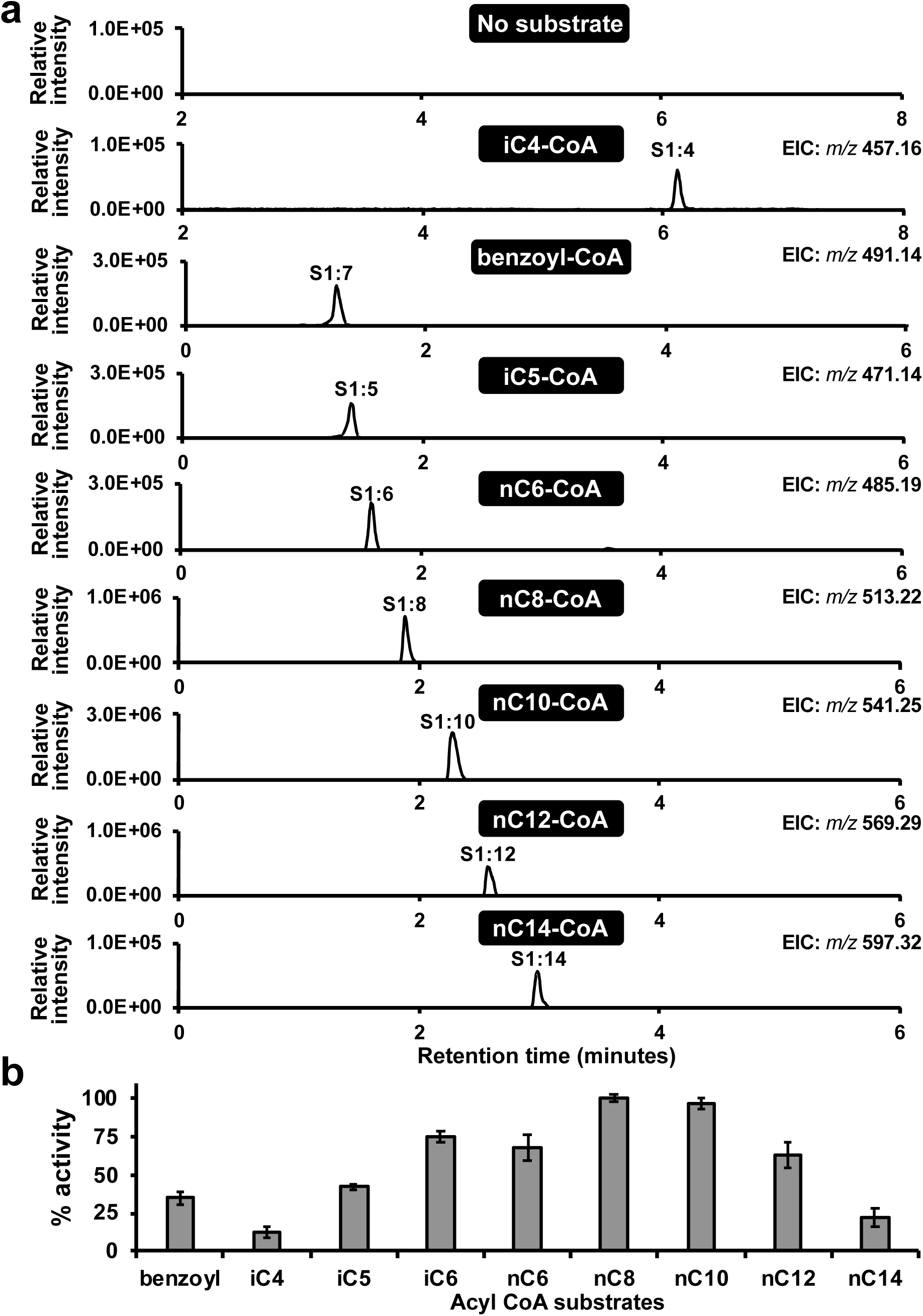
*In vitro* activity and acyl acceptor promiscuity of NacASAT1. **(a)** *In vitro* enzyme assays were conducted with NacASAT1 together with 1 mM sucrose and 250 μM CoA. Assays were analyzed using LC-MS and extracted ion chromatograms (EIC) are shown for the corresponding monoacylated sucrose products. All assay products were analyzed on a C18 column, except those produced with iC4-CoA, which was separated using a BEH amide column. **(b)** Quantification of assays in **(a)** were performed in triplicate and the area of the monoacylated product was corrected using an internal standard. Percent activity was set to 100% for the substrate with highest activity, nC8-CoA. Data shown are the mean of three biological replicates ± SEM.

The combined VIGS and enzyme assay results led us to test the acylation position of monoacylated products. Assays were performed with sucrose and iC5, nC6 or nC8 CoAs and the retention times of the monoacylated products were compared to those from ASAT1 enzymes with known acylation positions: 2-position for. *S. sinuata* SsASAT1 (Moghe et al., 2017) and 4- position for tomato SlASAT1 (Fan et al., 2016). The monoacylsucrose products of NacASAT1 and SsASAT1 with iC5-, nC6-, and nC8-CoA matched retention times using two different LC column chemistries, indicating that NacASAT1 acylated sucrose at the 2-position (Supplemental Figure S10). Acylation at the 2-position was observed for the NMR-resolved acylsugars, consistent with the hypothesis that acylation at this position is the first step in *N. acuminata* acylsugar assembly.

### NacASAT4 acts as an acetyltransferase that uses *in vivo* metabolites as substrate

In contrast to the first step of the pathway, it is more challenging to perform *in vitro* validation of NacASAT candidates for the intermediate and late-stage steps of acylsugar biosynthesis. This is because pathway intermediates such as mono, di and triacylated sucroses do not accumulate in *N. acuminata*, making the substrates of these reactions difficult to obtain. A notable exception was identified in the NMR-resolved tetra and pentaacylated sucrose structures from *N. acuminata*: these differ in acetylation, which facilitated identification of an acetyltransferase enzyme. We demonstrated activity of the NacASAT4 ortholog using the NMR-resolved *N. acuminata* S4:16(2,4,5d,5d) and acetyl-CoA substrates to produce S5:18, which had identical retention time as S5:18(2,2^R6′^,4,5d,5d) from *N. acuminata* plant extracts (Supplemental Figure S11a,b). This result is consistent with the hypothesis that NacASAT4 acetylates the 6′ position of tetraacylsucroses. To determine if NacASAT4 could acetylate triacylsucroses, we took advantage of the observation that acyltransferase enzymes catalyze the reverse reaction in the presence of free CoA (Leong et al., 2020; Lou et al., 2021); indeed NacASAT4 deacetylated S4:16(2^R3′^,4,5d,5d) at the 3′ position to produce an S3:14 product (Supplemental Figure S11c). These results are consistent with the hypothesis that NacASAT4 acts as an acetyltransferase that can perform consecutive reactions converting S3:14(4,5d,5d) into S5:18(2^R3′^,2^R6′^,4,5d,5d).

Although previously characterized ASAT4 acetyltransferases function late in acylsugar assembly (Fan et al., 2016; Moghe et al., 2017; Leong et al., 2020; Lou et al., 2021), we asked whether NacASAT4 could also act early in acylsugar assembly using sucrose or monoacylated sucrose as acceptor substrates. However, no products were observed in assays with acetyl-CoA, sucrose, and NacASAT4 (Supplemental Figure S12). We also tested whether acyl-CoAs other than C2 were suitable donor substrates. In assays containing NacASAT1 together with NacASAT4, nC6-CoA or nC8-CoA and sucrose only monoacylated products at the 2-position were observed (the result of NacASAT1 activity; Supplemental Figure S12). Taken together, these *in vitro* enzyme assay results are consistent with the hypothesis that NacASAT4 is an acetyltransferase acting late in the *N. acuminata* acylsugar pathway.

### *In vitro* synthesis of structurally diverse acylsugars using *Nicotiana* acyltransferases

The striking diversity of acylsugar variants found across *Nicotiana*, along with the acyl-CoA donor substrate promiscuity of NacASAT1, suggests that NacASATs can be useful tools for synthetic biology. To explore the *in vitro* capability of acyltransferases to make acylsugar products, NacASAT1, NacASAT4 and the other two orthologous acyltransferases identified through comparative transcriptomics, NacASAT2 and NacASAT3, were analyzed for activity in ‘one pot’ assays with sucrose as acceptor, with iC5- and C2-CoAs used as acyl donor substrates because these were commonly found in *N. acuminata* acylsugars (Supplemental Figure S1). This reaction produced seven detectable acylsucrose products ranging from mono to pentaacylated (Figure 5a). This result demonstrates the ability of NacASATs to make varied acylsugars *in vitro* with only two acyl CoA substrates (Figure 5).

**Figure 5.**
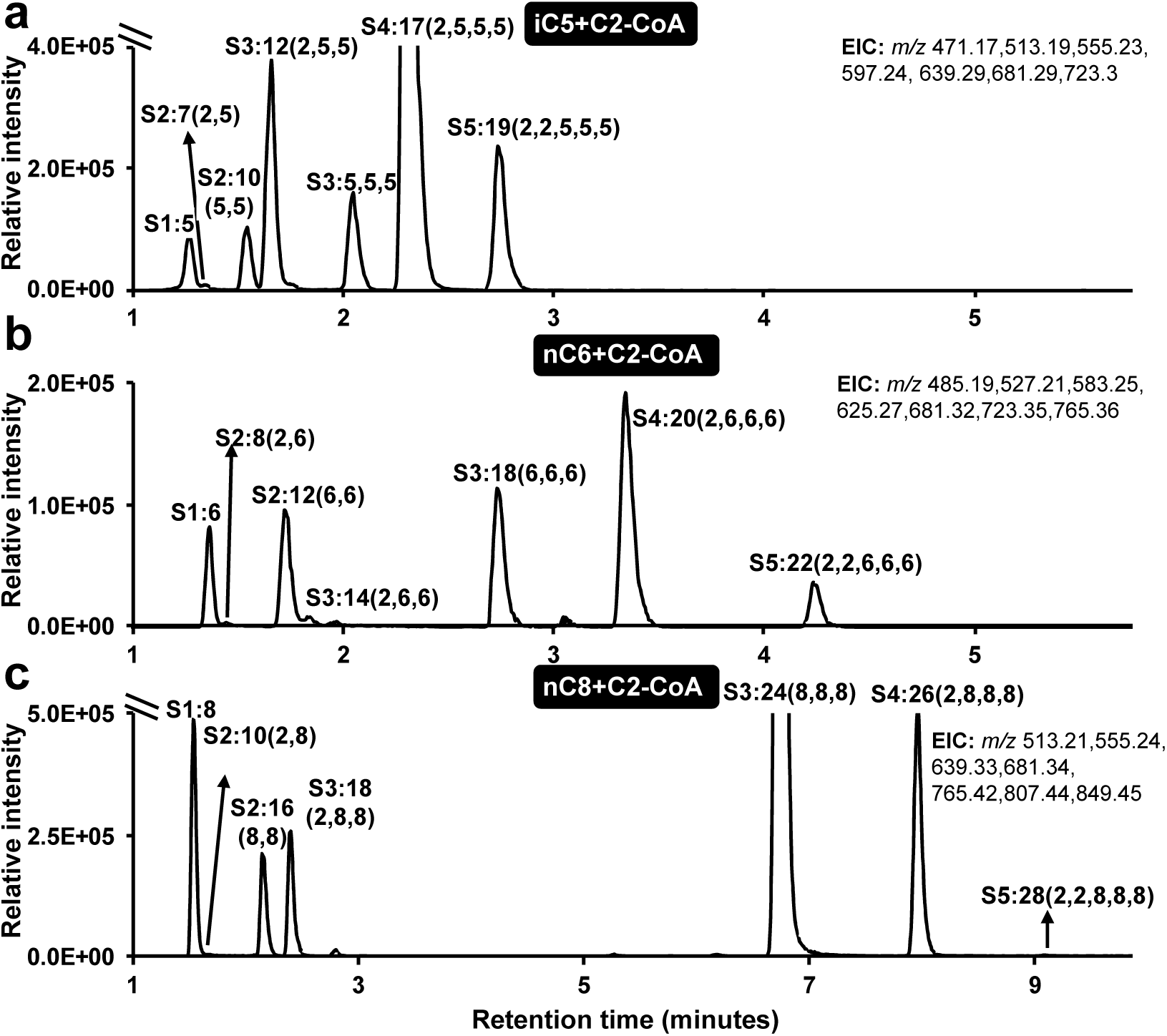
*In vitro* enzymatic production of acylsugar structural variants. *In vitro* assays containing 1 mM sucrose and 250 μM various indicated acyl CoAs were incubated with NacASATs1-4 for 30 minutes at 30°C and monitored using LC-MS for acylated sucrose products. Assays with iC5 and C2-CoA **(a)**, nC6 and C2-CoA **(b)**, or nC8 and C2-CoA **(c)**. Extracted ion chromatograms (EIC) are shown for the predicted masses of the corresponding acylsugars, *m/z* of the EIC are shown in the top right of each chromatogram. The number above each peak corresponds to the annotated acylsugar product based on accurate mass and fragmentation patterns. Note that the y-axis is truncated in some chromatograms to show low accumulating products.

Given the ability of NacASAT1-4 to produce multiple acylsugar products using only two acyl CoA substrates, we tested their activities with CoA donor substrates corresponding to those less commonly found on *Nicotiana* acylsugars. Seven *in vitro* products each were detected when either nC6- or nC8-CoA were used with C2-CoA, sucrose and NacASAT1-4 (Figure 5b & c). In assays with permutations containing C2-CoA with nC6- or nC8-CoA, products were detected that contain one to five acylations, demonstrating the capacity of these enzymes to produce varied acylsugar structures *in vitro*. While the relative quantities of products depended on the CoA substrate combinations used (Figure 5), the more highly acylated products generally produced a stronger signal than the less acylated products, which may reflect differing ionization efficiencies of the products (Ghosh and Jones, 2015). For example, in assays with iC5 and nC6 CoA the predominant products were the tetraacyl sucroses S4:17(2,5,5,5) and S4:20(2,6,6,6), respectively (Figure 5a & b), whereas in assays with nC8-CoA the triacyl sucrose S3:24(8,8,8) predominated (Figure 5c). Positive-ion mode analysis of diacetylated *in vitro* products such as S5:22(2^P^,2^F^,6,6,6) revealed one acetylation on each ring (Supplemental Figure S13). This contrasts with the pentaacylated acylsugars extracted from *N. acuminata*, which have both acetylations on the furanose ring (Figure 2c). In contrast to the large variety of products produced with iC5, nC6 and nC8-CoA, a one pot reaction with C2-CoA and iC4-CoA, NacASAT1-4 and sucrose led to detection of only S1:4 (Supplemental Figure S14a). Taken together, one pot acylsugar assays with various acyl-CoA substrates found on *in planta* derived acylsugars demonstrates the synthetic capacity of NacASAT1-4 to make many acylsugar products. Furthermore, most of the products that accumulated in these assays were not detectable from *N. acuminata* plant extracts, demonstrating that there are additional factors driving acylsugar chemical diversity *in planta*.

To further assess acyl-CoA donor promiscuity and attempt to enhance the acylsugar product output from *in vitro* assays, NacASATs were tested in one pot assays with acyl-CoA substrates not detected in the *in planta*-produced acylsugars across the *Nicotiana* genus, including longer (nC10-nC14) and aromatic (benzoyl) acyl-CoAs. Five acylsugar products were detected using nC10- and C2-CoA: S1:10, S2:12(2,10), S2:20(10,10), S3:22(2,10,10), and S3:30 (10,10,10) (Supplemental Figure S14c). This trend towards less complex product mixtures increased with the longer donor substrates: only S1:12(12) and S2:14(2,12) or S1:14 (14), and S2:16(2,14) products were observed in assays containing C2-CoA and nC12 or C14, respectively (Supplemental Figure S14d & e). In assays with benzoyl-CoA (C7^B^-CoA), C2-CoA, sucrose and NacASAT1-4, only the monoacylated sucrose product S1:7^B^ was observed (Supplemental Figure S14b), but no further acylated products, similar to the product profile of NacASAT1 tested with single CoA substrates (Figure 4). These assays further highlight the catalytic promiscuity of NacASAT1, but reveal that subsequent acylating enzymes may be more specific and cannot accommodate bulkier acylated sucroses.

The *in vitro* product diversity of one-pot assays using two different acyl-CoA substrates at a time led us to explore how much acylsucrose complexity could be produced by including larger numbers of acyl-CoA substrates with NacASATs. NacASAT1-4 were tested together in a reaction with a mixture of sucrose and equimolar C2, nC6, nC8 and nC10 CoAs. Over 25 products were detected based on chromatographic retention, accurate mass, and fragmentation patterns, exceeding the acylsugar complexity of *N. acuminata* surface metabolite extracts (Table 1, Supplemental Table S1 & Supplemental Figure S15). Inclusion of non-natural acyl-CoA substrates in these assays led to production of acylsugars that were not identified in plant extracts. Because of the similar retention times for many of the acylsugar products, the chromatograms were individually searched using the predicted accurate mass of the products. As documented in Table 1 and Supplemental Figure S15 three monoacylated and eight diacylated products were identified. The latter could be classified into two types: ones with an acetylation plus a long acyl chain (e.g. S2:8(2,6)) and those with two long acyl chains (e.g. S2:12(6,6)). The monoacylated and the acetylated diacylsugars produced in this assay were not detected in any *Nicotiana* species screened (Table 1, Supplemental Table S1 & Supplemental Figure S15). Eleven triacylated products were detected, which could be grouped into two classes; those with an acetylation and those without an acetylation (Table 1 & Supplemental Figure S15). Some products could not be differentiated because they have the same accurate mass, were not well separated and did not accumulate to a sufficient level that fragmentation patterns could distinguish the two isomers (for example the structural isomers S4:26(2,8,8,8) and S4:26(2,6,8,10); Supplemental Figure S15). Unlike assays that contained only two CoAs (Figure 5), no pentaacylated products were observed in this mixed acyl donor assay (Supplemental Figure S15). Because of the inclusion of nC10-CoA, many of the products from these assays were not detected in any *Nicotiana* plant extracts (Table 1), further demonstrating the metabolic potential of NacASATs using an *in vitro* acylsugar production platform.

**Table 1.**
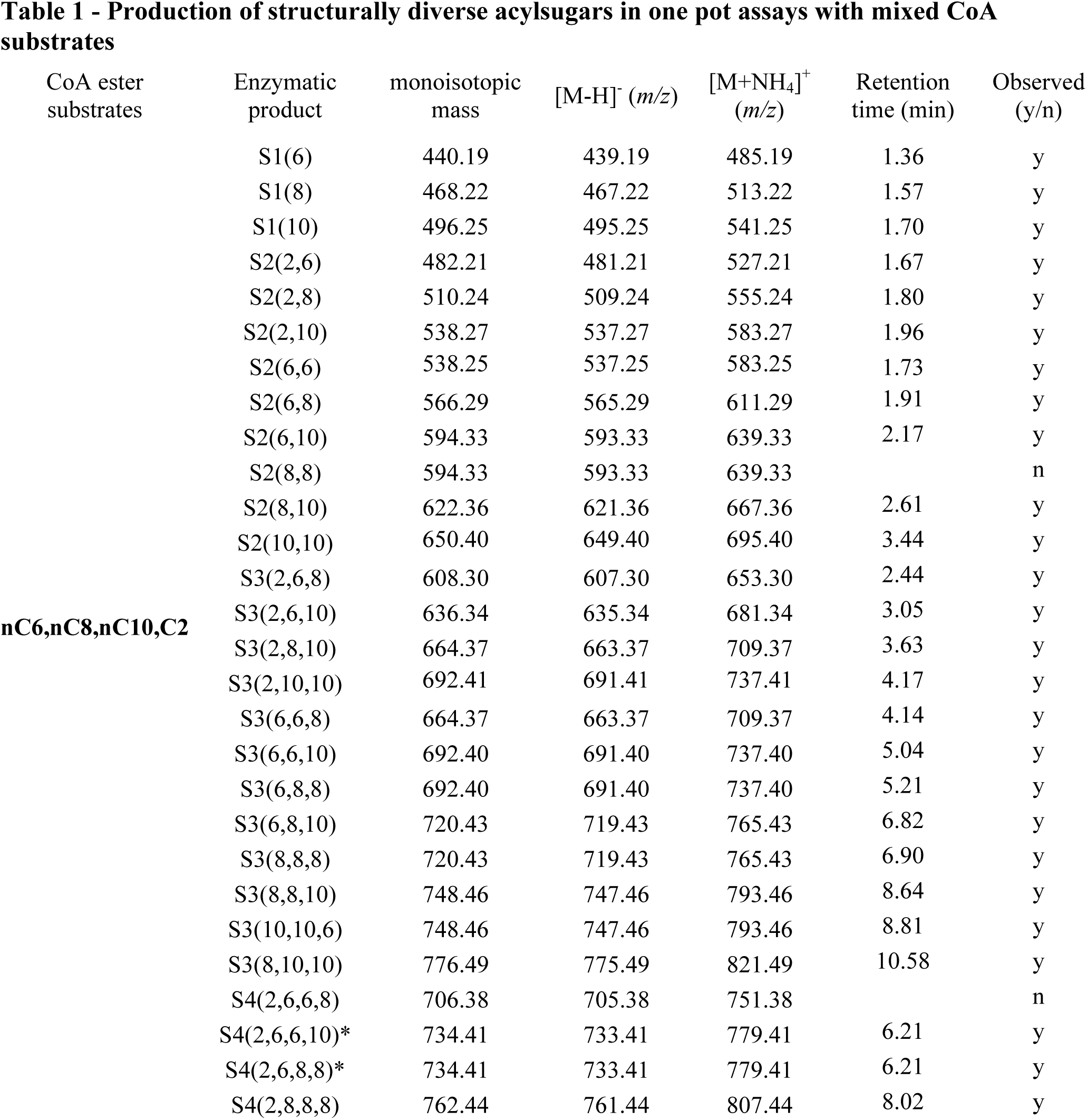
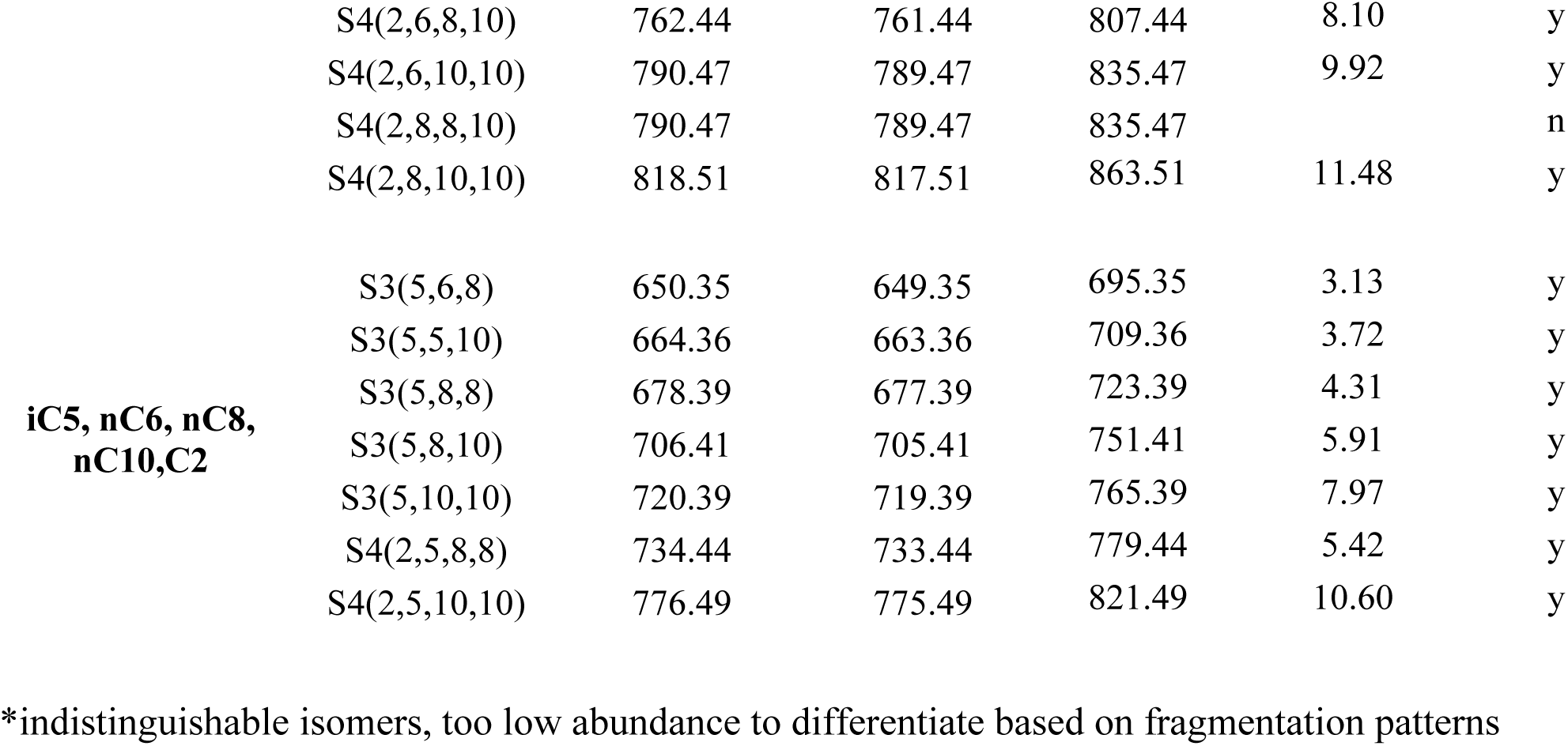
Production of structurally diverse acylsugars in one pot assays with mixed CoA substrates

The structurally varied products produced from the above assay led us to ask if we had reached the upper limit of enzymatic capacity in the *in vitro* assays. To address this, we added an additional acyl-CoA substrate into the *in vitro* assay. Because iC5-CoA was used by NacASAT1- 4 to produce various acylsugars *in vitro* (Figure 5), iC5-CoA was combined with equal concentrations of C2, nC6, nC8, and nC10 CoAs and 1 mM sucrose in assays containing NacASAT1-4. In addition to the acylsugars produced in the mixed-CoA assay with C2, nC6, nC8, and nC10 CoAs (Supplemental Figure S15), seven other structural variants were produced with iC5 chains (Table 1, Supplemental Figure S16). For example, a few of the more abundant products were S3:21(5,8,8), S3:23(5,8,10), and S4:27(2,5,10,10) enhancing the number of acylsugars produced *in vitro* in a single assay (Table 1, Supplemental Figure S16).

### Cross-species ASAT mixed assays highlight promiscuity and regiospecificity of ASAT1s

The array of products produced in one pot reactions led us to ask whether additional products could be assembled by combining ASATs from different Solanaceae lineages. The observation that SlASAT1 and NacASAT1 acylated at different positions provided an opportunity to test if two divergent ASAT1s could produce diacylsucroses. We tested downstream NacASAT enzymes with *S. sinuata* ASAT1 (SsASAT1) and tomato ASAT1 (SlASAT1), which acylate at the 2- and 4- positions, respectively (Supplemental Figure S10; Fan et al., 2016; Moghe et al., 2017). Only monoacylated products were detected when the 4-position acylating SlASAT1 was combined with NacASAT2-4, C2 and nC10-CoA substrates, (Supplemental Figure S17), demonstrating that NacASAT2-3 did not use 4-position acylated monoacylated sucroses as substrates. In contrast, SsASAT1 and NacASAT2-4 together with C2- and nC10-CoA and sucrose produced the same products as assays with NacASAT1-4 (Supplemental Figure S17), albeit in different relative abundances. These data support that SsASAT1 and NacASAT1 perform the same function *in vitro* (Supplemental Figure S17), acylation of sucrose at the 2-position.

SlASAT1 and NacASAT1 were combined and tested with sucrose and nC6-CoA as both enzymes used these substrates. This assay produced both diacylated (S2:12) and monoacylated S1:6 products (Supplemental Figures S18a & 19a), consistent with the hypothesis that R2 and R4 acylating ASAT1s together can produce diacylsugars. To test the order of acylation producing S2:12, we performed sequential assays with nC6-CoA and sucrose in which NacASAT1 incubation was followed by heat inactivation, and then addition of SlASAT1, or the assay was performed in the opposite direction. In the sequential assay with NacASAT1 added first, both mono and diacylated products were observed (Supplemental Figure S18b). In contrast, when SlASAT1 was added first, only mono-acylated products were produced and multiple peaks were identified, likely due to the previously reported (Fan et al., 2016; Leong et al., 2019; Lou et al., 2021) heat and base-promoted non-enzymatic rearrangement of the 4-position monoacyl product (Supplemental Figure S18b). These assays suggest that SlASAT1, but not NacASAT1, can acylate S1:6^R2^ *in vitro*. To test this hypothesis, purified S1:10(nC10) acylated at the 2- or 4- position (Lou et al., 2021) were used as acyl acceptor substrates with nC6-CoA as the acyl donor in assays together with NacASAT1 or SlASAT1. SlASAT1 converted S1:10^R2^ into S2:16(6,10^R2^) but did not transform 4-position acylated sucrose into a diacylsucrose (Supplemental Figure S19b). In contrast, NacASAT1 was unable to use either 2- or 4-position monoacylated sucroses as substrates (Supplemental Figure 19b). These results demonstrate that SlASAT1 can acylate both sucrose and 2-position monoacylated sucrose while NacASAT1 only acylated nonesterified sucrose. The acyl acceptor promiscuity of SlASAT1 and the acyl donor promiscuity and acceptor regiospecificity observed for NacASATs highlight the varied properties and activities of ASATs from across the Solanaceae family. This work demonstrates the utility of using acyltransferases in synthetic biology platforms for engineering structurally diverse acylsugars for optimal biological activity.

## DISCUSSION

The evolution of plant specialized metabolism is potentiated by distinct incremental changes that together profoundly impact plant chemical diversity. The varied activities of lineage-specific specialized metabolic enzymes convert relatively simple core metabolite precursors – sugars and acyl-CoA’s for acylsugars – into an array of specialized metabolites. Understanding mechanisms that contribute to metabolic innovations can lead to new synthetic biology tools and strategies, which then can be harnessed to expand upon the structures of biologically produced active metabolites. Here, we expanded upon acylsugars produced across the *Nicotiana* genus using *in vitro* enzymatic synthesis of non-natural acylsugar products.

### Acyl chain variation and distribution across the family

Screening acylsugars across the *Nicotiana* revealed over 300 annotated acylsugar structural variants (Supplemental Table S1). These differences are due to accumulation of glucose and sucrose esters (Supplemental Table S1, Supplemental Figure S5), as well as varied types, ester positions, and numbers of acyl chains attached to the sugar core (Figures 1, 2 & Supplemental Figures S1, S6). Acylsugar diversity placed in a phylogenetic context revealed that some acyl chains were detected in nearly all *Nicotiana* species screened: two such examples are iso and anteiso-C5 acyl chains, derived from leucine and isoleucine, respectively (Figure S1; Binder, 2010). C5-CoA acyl chains are commonly found on acylsugars across the Solanaceae (Kroumova et al., 2016; Moghe et al., 2017), suggesting that ASATs capable of using C5-CoAs and the upstream metabolic pathway providing C5-CoA substrates are highly conserved. In contrast, phylogenetic restriction of some acyl chain types highlights potential metabolic innovations potentiating acylsugar structural variation. For example, C2 acyl chains are absent in a single *Nicotiana* clade, which suggests the possibility of *ASAT4* gene inactivation or loss in an ancestor. This is analogous to variation in acetylated acylsugars across *S. habrochaites* accessions, which correlated with gene mutation of *ShASAT4* (*AT2*) in accessions lacking acetylated acylsugars (Kim et al., 2012). Whereas there is evidence that *ASAT4* gene mutation occurred multiple times in *S. habrochaites*, the most parsimonious explanation for lack of C2 acylations in one *Nicotiana* clade is a single loss or inactivation event. Another example was the phylogenetic restriction of tiglylated acylsugars (Figure 2 & Supplemental Figures S1, S6), suggesting a metabolic alteration in BCAA catabolism increasing substrate availability and enabling incorporation into acylsugar biosynthesis. Tiglyl acyl chains were detected within a single *Nicotiana* clade, and although present in other specialized metabolites such as tropane alkaloids (Leete, 1973; Rabot et al., 1995), are uncommon in acylsugars (Matsuzaki et al., 1991; Zhang et al., 2016). These results demonstrate how specific and distinct changes in metabolic pathways can have major impacts on specialized metabolite structures across a species or genus.

Past studies suggested that acyl chain length is another phylogenetically restricted trait, and indeed we detected no acyl chains > 8 carbons long in *Nicotiana* (Supplemental Figure S1). The lack of long acyl chains in *Nicotiana* is also found in closely related species (Kroumova et al., 2016; Moghe et al., 2017; Nadakuduti et al., 2017), but distinct from plants in the Solanaceae genera such as *Physalis*, *Solanum* and *Hyoscyamus*, which accumulate acyl chains up to 14 carbons long (Ghosh et al., 2017; Zhang et al., 2016; Moghe et al., 2017). The phylogenetic distribution of acylsugar acyl chain lengths observed could result from differences in ASAT specificity and alterations in upstream metabolic pathways leading to variations in trichome CoA availability. The data reported here demonstrate that NacASATs can incorporate structurally diverse CoAs with a preference for longer chain CoAs (Figures. 4, 5 & Supplemental Figure S14), which do not accumulate on *in planta* acylsugars; this is consistent with the hypothesis that trichome CoA availability contributes to acyl chain phylogenetic distribution. Manipulation of acyl chain elongation pathways through transgenic approaches to alter trichome CoA pools may be another way to engineer structural variants of acylated natural products in plants and synthetic production systems.

### Synthesis of *in vitro* acylsugar products not detected in *Nicotiana* trichomes

Our documentation of *in vitro* products not detected in surface extracts of any of the *Nicotiana* species tested has implications for understanding BAHD structure-function relationships, and engineering novel acylated natural products. Multiple nC10-, nC12-, and nC14-containing acylsugars accumulated *in vitro*, but were not found in any *Nicotiana* species analyzed (Supplemental Figures S14, S15 & S16; Table 1 & Supplemental Table S1). The observed *in vitro* promiscuity of ASATs in one pot assays with multiple acyl-CoA substrates further highlights *in vivo* metabolic constraints limiting acylsugar structures produced by plants.

By taking advantage of enzymatic activities that link upstream core metabolism to specialized metabolic pathways, synthetic biology platforms can further increase the structural outputs when coupled with promiscuous ASATs. These assays demonstrate the possibility of using an *in vitro* synthetic biology platform, along with transgenic modification, to expand upon the numbers of metabolites produced *in planta*.

### Glucose and sucrose core distribution

Our results extend previously published work demonstrating the presence of acylsugars based on both glucose and sucrose cores across the *Nicotiana* (Matsuzaki et al., 1989, 1991; Shinozaki et al., 1991; Figure S5). Acylsucroses are found across the Solanaceae family (Moghe et al., 2017), whereas acylglucoses are less common and sporadically distributed; for example acylglucoses are found in phylogenetically distinct *Solanum* species (Leong et al., 2019; Lou et al., 2021). In *Solanum*, acylglucoses are derived from acylsucroses by the activity of acylsucrose fructofuranosidase (ASFF) enzymes that cleave the glycosidic linkage (Leong et al., 2019; Lou et al., 2021). Variation in sugar core composition was observed across geographically distinct *S. pennellii* lineages (Shapiro et al., 1994; Lybrand et al., 2020). Because of the sporadic distribution of acylglucoses across the *Nicotiana* genus (Supplemental Figure S5), it is unclear if this trait independently evolved multiple times as seen in *Solanum* (Leong et al., 2019; Lou et al., 2021). Production of acylsugar mixtures consisting of various sugar cores and acyl chains both *in vitro* and *in planta* (Table 1 & Supplemental Table S1) provides an opportunity to investigate their biological significance. For example, mixtures of acylsugar variants were shown to possess more potent antiherbivore activity compared to a single acylsugar structure (Puterka et al., 2003), suggesting that broadening acylsugar types through metabolic engineering or biochemistry-informed breeding may enhance plant resilience.

## Materials and Methods

### Plant growth

*Nicotiana* seeds were obtained from the U.S. Department of Agriculture Germplasm Resources Information Network (GRIN; https://www.ars-grin.gov/). Seeds were treated with a solution of 10% (w/v) trisodium phosphate and 10% bleach for 5 minutes with constant rocking. Seeds were rinsed six times with distilled water, placed on Whatman filter paper and germinated at 28°C in the dark. Seedlings were transplanted into peat pots (Jiffy) and placed in 3” square pots and grown at 24°C under a 16/8-hour day/night cycle with ∼70 μmol m^-2^ s^-2^ photosynthetic photon flux density using cool white fluorescent bulbs. Plants were fertilized once per week with half strength Hoagland solution.

### Acylsugar extraction, quantification and detection

Acylsugars used for sugar core analysis were extracted from ∼6-week-old plants from fully expanded leaves. Intact leaves were placed in a glass tube with 2 mL of H_2_O and 4 mL of dichloromethane and gently vortexed for 30 seconds. Following phase separation, 1 mL of the organic phase was transferred to a glass tube and dried to completeness under vacuum, then dissolved in 1 mL of isopropanol + 0.1% formic acid and vortexed for 30 seconds.

For acylsugar core analysis, 20 μL of the acylsugar extract was added to 200 μL of methanol and 200 μL of 3 M ammonium hydroxide in a 1.5 mL microfuge tube. Samples were vortexed for 20 seconds and placed at 22°C for 48 hours. The samples were briefly centrifuged and dried under vacuum at 35°C. The pellet was dissolved in 200 μL of 10 mM ammonium bicarbonate pH 8.0 in 80% acetonitrile containing the isotopically labeled quantitation standards 1 μM [^13^C_12_]sucrose and 1 μM [^13^C_6_]glucose and analyzed using a Waters Acquity TQ-D UPLC-MS-MS equipped with a Waters Acquity UPLC BEH amide column (2.1×100 mm, 1.7 μm) as previously described (Lybrand et al., 2020).

For acylsugar identification using a Waters Acquity UHPLC coupled to a Waters Xevo G2-XS QToF mass spectrometer, acylsugars were extracted from a single fully expanded leaf from ∼6-week-old plants. A leaf was placed into 1mL of extraction solvent (3:3:2 acetonitrile:isopropanol:water + 0.1% formic acid + either 10 μM propyl 4-hydroxybenzoate or 1 μM telmisartan internal standard) and gently mixed for 2 minutes. Acylsugar extracts (10 μL) were run on an Ascentis Express C18 HPLC column (100 mm x 2.1 mm, 2.7 μm) as described previously (Leong et al., 2019; Fan et al., 2020; Lou et al., 2021). Differences in LC gradients and mass spectrometer parameters are noted below and in Supplemental Table S2.

Acylsugars were annotated through collision induced dissociation (CID) at increasing collision cell potentials based on fragmentation patterns in ESI^-^ and ESI^+^ modes. For ESI^-^ mode the following mass spectrometer settings were used. Three acquisition functions were used at quasi-simultaneous stepped collision cell potentials (0, 20, 45 V) to acquire spectra. The source temperature was 100°C, the desolvation temperature was 350°C, capillary voltage was 2 kV, cone voltage was 40 V, the mass range collected was *m/z* 50 to 1,500. Lock mass correction using Leu-enkephalin [M-H]^-^ as the reference was applied during data collection. For ESI^+^ mode the following mass spectrometer settings were used. Three acquisition functions were used at quasi-simultaneous stepped collision potentials (0, 25, 50 V) to acquire spectra. The source temperature was 100°C, the desolvation temperature was 350°C, capillary voltage was 2 kV, cone voltage was 40 V, the mass range observed was *m/z* 50 to 1,500. Lock mass correction using Leu-enkephalin [M+H]^+^ as the reference was applied during data collection.

Acylsugar acyl chain analysis was performed as previously published (Fan et al., 2020, Ning et al., 2015). Acyl chains were determined through library matches of the mass spectra of the corresponding ethyl esters using NIST Library Version 14. The relative abundances were calculated by integrating the corresponding peak area based on total ion current (TIC) over the total acyl chain peak area.

Sugar core composition and acyl chain types were placed into a taxonomic context based on various phylogenetic analyses of the *Nicotiana* genus (Bally et al., 2018; Clarkson et al., 2004; Clarkson et al., 2017; Särkinen et al., 2013).

### LC QTOF-MS annotation of acylsugars

Acylsugar structures were inferred from experimentally determined exact masses and fragmentation patterns in positive and negative ion modes. Two discrete collision energies in negative ion mode were used, the lower energy was used to generate fragment ions corresponding to the loss of the acyl chains and the higher energy created carboxylate ions of the corresponding acyl chain losses. Positive ion mode fragmentation patterns were used to infer positions of acetyl groups on acylsugars, as cleavage of the glycosidic linkage typically occurs prior to ester bond cleavage. Acylsugars were manually annotated as demonstrated in Supplemental Figures S2 and S3. Acylsugars with similar retention times (+/- 0.05 minutes), exact masses (+/- 10 ppm) and fragmentation patterns were considered to be the same acylsugar and highlighted in yellow in Supplemental Table S1.

### Acylsugar extraction, purification, and NMR analysis

For bulk acylsugar extraction, aerial tissue was harvested from six ∼6-week-old plants in 1 L of extraction solvent (1:1 acetonitrile:isopropanol + 0.1% formic acid). Plant material was swirled in a large flask for two minutes and then the extract was filtered through a single layer of Whatman filter paper. A rotary evaporator was used to dry the extract to completeness. The dried extract was dissolved in 15 mL of extraction solvent and stored at -80°C. A Waters 2795 preparative HPLC equipped with a Dionex Acclaim C18 column (4.6 x 150 mm, 5 μm, 120 Å) was used for fractionation and purification of isolated acylsugars. Twenty injections of the acylsugar extract (100 μL each) were applied onto the HPLC using solvents A (acetonitrile) and B (water + 0.1% formic acid) and fractions were collected every 20 s. The 60-minute acylsugar screening gradient (Supplemental Table S2) was used to separate closely eluting acylsugars. Fractions were checked for purity by running on a Waters Xevo G2-XS QToF mass spectrometer as described above and fractions were pooled that contained isolated acylsugars. ∼1.0 mg of purified compound was dissolved in 300 µL of deuterated NMR solvent CDCl_3_ (99.96% atom % D, Sigma-Aldrich) and transferred to solvent-matched Shigemi tube for analysis. ^1^H, ^13^C, COSY, HSQC, HMBC, TOCSY and NOESY NMR experiments were performed at the Max T. Rogers NMR Facility at Michigan State University using a Bruker Avance 900 spectrometer equipped with a TCI triple resonance cryoprobe. All spectra were referenced to non-deuterated chloroform solvent signals (δH = 7.26 (s) and δC = 77.2 (t) ppm). NMR spectra were processed using MestReNova 12.0.1 software.

### RNA extraction, quantification, library preparation, and RNA sequencing

*N. acuminata* plants were grown until flowering, and stem tissue was used for trichome isolation. Stem tissue was frozen in liquid N_2_ and the trichomes scraped as reported previously (Ning et al., 2015; Fan et al., 2016; Moghe et al., 2017; Nadakuduti et al., 2017). Three plants were used for trichome isolation for each biological replicate RNA extraction. RNA was extracted from isolated trichomes and the underlying shaved stem tissue using a Qiagen RNAeasy Plant Mini Kit (Qiagen) following the kit protocol with an on-column DNaseQ treatment (Qiagen). RNA was quantified using a Qubit fluorescence assay following the kit protocol. Three biological replicates for each tissue (shaved stem and trichome) were used for library preparations. Libraries were prepared using the Illumina TruSeq Stranded mRNA Library Preparation Kit on a Perkin Elmer Sciclone G3 robot following manufacturer’s recommendations. Libraries were pooled and loaded in equimolar concentrations onto a single lane of an Illumina HiSeq 4000 flow cell and sequencing was performed in a 2×150bp paired end format using HiSeq 4000 SBS reagents. Base calling was done by Illumina Real Time Analysis (RTA) v2.7.7 and output of RTA was demultiplexed and converted to FastQ format with Illumina Bcl2fastq v2.19.1. Library prep and sequencing was performed by the RTSF Genomics Core at Michigan State University

### Transcriptome assembly and quality assessment

RNA-seq reads (submitted to the sequence read archive (SRA): PRJNA740342) were analyzed for quality using FastQC. RNA-seq reads were then trimmed using Trimmomatic v0.38 with the following parameters ILLUMINACLIP:TruSeq3-PE.fa:2:30:10 LEADING:3 TRAILING:3 SLIDINGWINDOW:4:15 MINLEN:75, all other parameters were set as default. The trimmed reads were used for *de novo* transcriptome assembly using Trinity v2.6.6 with the following parameter min_contig_length 350, all others were set as default. A total of 174,800 transcripts were detected and collapsed into 65,168 trinity ‘genes’ based on the longest isoform. The average contig length was 1333 nucleotides with an N50 value of 2048 nucleotides. Reads were then mapped to the *de novo* assembly and counted using Bowtie v1.2.2 and RSEM v1.3.1with default parameters. An MDS plot of the biological replicates indicates that the trichome samples cluster together and are distinct from the shaved stem tissues samples (Supplemental Figure S20). The longest open reading frames for each transcript were identified and translated using TransDecoder v2.1.0 with the default settings. Orthologous groups were then predicted from the predicted protein sequences of *N. acuminata* together with protein sequences from *S. lycopersicum*, *S. pennellii*, *S. tuberosum*, *N. tabacum*, and *A. thaliana* using OrthoFinder v2.2.7 with default settings.

### VIGS Analysis

VIGS fragments targeting two different regions of *NacASAT1* (sequence 5’-3’ ASAT1a, aaatccaactcgggtagaaatactcacagcacttttttataaatgtggtatgaaagtgatcaattcgtgttcattcaagccatctatcttattcctaactgtgaatttaagatctcttattcctctgccagatgatactcctggaaattttagttcttcccttcttgtgcctacatataatgaagaagaaatgaatttatcaagattggttagtcagctaa; ASAT1b, tgctgattttttcaaggctcgattcgattgtcccatgtctgaaatccttaaaagtcctgataaaaatgtcaaagaattagtatatcctaaggatataccatggaatgttgttacatctaatagaaagttggtcacagttcaatttaaccaatttgattgtggaggaatagctctaagtgcatgtgtatcacacaaaattggagatatgtgcacagtatccaaatttttacaag) and *NacPDS* (sequence 5’-3’ ggagggcaatcttatgttgaagctcaagacggtttaagtgttaaggactggatgagaaagcaaggtgtgcctgatagggtgacagatgaggtgttcattgccatgtcaaaggcacttaacttcataaaccctgacgagctttcaatgcagtgcattttgattgctttgaacagatttcttcaggagaaacatggttcaaaaatggcctttttagatggtaaccct) were designed using SGN VIGS tool (https://vigs.solgenomics.net/) and the fragments were amplified using PCR adapters for ligation into pTRV2-LIC. pTRV2-LIC was digested as previously described using *Pst*I to generate the linearized vector (Dong et al., 2007). Linearized vector and PCR fragments were combined using ligation independent cloning (LIC, NEB). Reactions were transformed into chemically competent *E. coli* Stellar cells. Following sequencing of isolated plasmids to confirm the correct sequence, plasmids including pTRV1 were transformed into *Agrobacterium tumefaciens* strain GV3101.

The VIGS protocol from Saedler and Baldwin 2004, which was used for gene silencing in *N. alata,* was adapted for *N. acuminata*. Following germination of *N. acuminata* seeds, once cotyledons emerged, seedlings were transferred into peat pots (Jiffy) and grown for 10 days at conditions stated above. Two days preinoculation, LB cultures containing kanamycin, rifampicin, and gentamicin were inoculated with pTRV1, pTRV2-NacASAT1, pTRV2-NacPDS, and an empty vector control (pTRV2-empty). Cultures were grown at 30°C at 225 rpm overnight in LB. Larger cultures composed of induction media (4.88 g MES, 2.5 g glucose, 0.12 g sodium phosphate monobasic monohydrate in 500 mL, pH 5.6, 200 μM acetosyringone), were inoculated using a 25:1 dilution of the overnight culture (50 mL total) and grown overnight grown at 30°C at 225 rpm. Cells were harvested by centrifugation at 3,200*g* for 10 minutes and gently resuspended in 1 volume of 10 mM MES, pH 5.6, 10 mM MgCl_2_. Following centrifugation at 3,200*g* for 10 minutes, cell pellets were resuspended in 10 mL of 10 mM MES, pH 5.6, 10 mM MgCl_2_. Cell suspensions were diluted using the same buffer to an OD600 of 1. Acetosyringone was added to the pTRV1 cell suspension to a final concentration of 400 μM. The different pTRV2-LIC constructs were mixed with an equal volume of pTRV1 suspension, resulting in a final acetosyringone concentration of 200 μM. Individual plants were inoculated through the abaxial side of the cotyledons and true leaves, if present. Plants were placed in the dark for 48 hours at 24°C and then returned to 16/8h day-night cycles at the same temperature. 14 days later, the plants were sampled for acylsugars and RNA using a bisected leaf for each experiment, typically the 4^th^ and 5^th^ true leaves, selected based on the *NacPDS* control plant photobleaching phenotype.

Acylsugars and RNA were extracted as described above. Acylsugars were quantified using Quanlynx software of an extracted ion chromatogram of individual acylsugars, corrected to the internal standard and expressed as the total acylsugar levels per dry weight of the leaf tissue. 10-15 biological replicates were used per VIGS fragment and T-tests were performed to determine if acylsugar levels were significantly different between empty pTRV2 controls and two independent pTRV2-NacASAT1 lines.

### qPCR analysis

RNA was extracted as described above from lines inoculated with NacASAT1 and a pTRV2- empty vector control. cDNA was synthesized using 1 μg of RNA using SuperScript II Reverse Transcriptase kit (Invitrogen) following the manufacturer’s directions. cDNA was diluted 10- fold following synthesis and pooled to determine the efficiencies of primers targeting *NacASAT1* (DN5384_c0_g1_i1), *NacEF1α* (DN11765_c1_g1_i7) and *NacActin2* (DN10810_c2_g1_i3). All primer efficiencies were between 90-110%. Primer sequences used for qPCR analysis can be found in Table S3. For qPCR reactions, cDNA was diluted a further 2-fold and 1 μL of cDNA was used as template in a 10 μL reaction with 1x SYBR Green PCR Master Mix (Thermo) and 200 nM forward and reverse primers. Reactions were carried out with a QuantStudio 7 Flex Real-Time PCR system (Applied Biosystems) by the RTSF Genomics Core at Michigan State University. The temperature cycling conditions were as follows: 50°C for 2 minutes, 95°C for 10 minutes, and 40 cycles of 95°C for 15 seconds and 60°C for 1 minute. Relative expression of *NacASAT1* was calculated using the ΔΔcT method (Livak and Schmittgen, 2001) and normalized to *NacEF1α* and *NacActin2*. 10-15 biological replicates were used with ≥ 3 technical reps used for the analysis, T-tests were performed to determine if *NacASAT1* transcript levels were significantly different between empty pTRV2 controls and two independent pTRV2-NacASAT1 lines.

### ASAT identification, cloning, expression, and protein purification

ASAT gene candidates from the *N. acuminata* transcriptome assembly were identified through BLAST searches using biochemically characterized ASATs from *S. lycopersicum*, *S. sinuata*, and *P. axillaris* (Fan et al., 2017, Moghe et al., 2017, Nadakauthi et al., 2018). Amino acid sequence alignments were performed using MUSCLE with default settings. All positions from the amino acid alignments with > 70% site coverage were used to build the phylogenetic trees in MEGA7 (Kumar et al., 2016) using maximum likelihood and 1,000 bootstrap replicates. Bootstrap values less than 50% were not shown. The tree was annotated using the interactive tree of life (iTOL; Letunic and Bork, 2019).

Primers for cloning were designed to amplify full length open reading frames and genes were amplified from cDNA prepared from *N. acuminata* leaf RNA and the primer sequences can be found in Table S3. Following gel purification using a Qiagen Gel Extraction Kit, open reading frames were cloned into a linearized pET28b vector with BamH1 and Xho1 in frame with a N-terminal 6x-His tag using Gibson HiFi assembly mix (NEB), 10 μL total reaction volume with 2 μL of linearized vector, 2 μL of insert, 5 μL of 2x HiFi mix and 1 μL of water. Reactions were incubated at 50°C for 30 minutes. For transformation into Stellar cells, 2 μL of the cloning mix was mixed with 50 μL of competent cells and placed on ice for 30 minutes. Following heat shock at 42°C for 60 seconds, cells were placed in a shaker at 37°C and 200 rpm with 1 mL of LB broth. Following sequencing to confirm the correct sequence, individual plasmids were transformed into BL21 Rosetta (DE3) protein expression cells (EMD Millipore).

For protein expression and purification, overnight cultures were added into 500 mL of LB broth with kanamycin (100 μg/mL) and chloramphenicol (20 μg/mL), when cultures reached an OD600 of 0.2, the incubator was turned down to 18°C and the shaking was reduced to 180 rpm. After 60 minutes isopropyl β-D-1-thiogalactopyranoside (IPTG) at 0.4 mM final concertation was added to induce protein expression and grown for additional 20 hours. The cultures were spun at 5,000*g* for 10 minutes and the pellet was dissolved in 25 mL of lysis buffer (25 mM HEPES pH 7.6, 50 mM NaCl, 10% v/v ethylene glycol + 0.5 M phenylmethylsulfonyl fluoride and 10 μM DTT). Cells were sonicated for 3 minutes then centrifuged for 30 minutes at 4°C at 50,000*g*. The supernatant was applied to a 1mL HisTrap FF column for purification of the recombinant protein using an Äkta Start FPLC system. The entire supernatant was loaded onto the column, then washed with 20 column volumes of Buffer A (0.5 M NaCl, 0.2 M sodium phosphate and 20 mM imidazole). The protein was eluted by increasing the concentration of Buffer B (0.5 M NaCl, 0.2 M sodium phosphate and 500 mM imidazole) and fractions judged by SDS-PAGE to contain the recombinant protein were pooled and desalted by size exclusion chromatography using 30K centrifugal spin column (Millipore).

### ASAT enzyme assays and detection of enzymatic products

Enzyme assays contained 250 μM acyl-CoA, 1 mM sugar core, 250 mM sodium phosphate buffer pH 6.0 and 10 μL of enzyme for a total of 30 μL, unless noted otherwise in the figure legends. Assays were incubated for 30 minutes at 30°C and stopped by addition of 60 μL of a mixture of 1:1 isopropanol and acetonitrile + 0.1% formic acid containing either 10 μM propyl- 4-hydroxybenzoate or 1 μM telmisartan internal standard. Reactions were placed on ice for 10 minutes and centrifuged at 17,000*g* for 5 minutes, the supernatant was transferred to a 2 mL glass LC vial with a 250 μL glass insert. Coupled assays were performed similarly to above. Enzyme assays with no CoA and with enzyme heated at 100°C for 10 minutes served as negative controls. Assays were run on LC-MS using either C18 and BEH amide columns with LC and mass spectrometer methods and settings similar to those used in acylsugar detection from plant extracts as described above. Acylsugar products were identified through accurate mass of expected acylsugar products and fragmentation patterns using extracted ion chromatograms.

## Acknowledgments

We thank the USDA Germplasm Resources Information Network (GRIN) for providing the *Nicotiana* seeds, the RTSF Genomics core, Mass Spectrometry and Metabolomics core and the Max T. Rogers NMR Facility at Michigan State University for sequencing, acylsugar and NMR analyses, respectively. Members of the Last lab including Rachel Kerwin for bioinformatics assistance. This research was funded by grants from the National Science Foundation to C.A.S. (IOS-1811055) and R.L.L. and A.D.J. (IOS-1546617). M.J. was a participant in the 2019 Plant Genomics at MSU Summer REU Program, supported by NSF grant DBI-1757043. A.D.J. acknowledges support from Michigan AgBioResearch through the USDA National Institute of Food and Agriculture, Hatch project number MICL02474.

## Data availability

*N. acuminata* ASAT amino acid sequences were deposited to NCBI: NacASAT1; MZ359196, NacASAT2; MZ359197, NacASAT3; MZ359198, NacASAT4; MZ359199. Raw reads from trichome and stem RNA-seq analysis were deposited to the Sequence Read Archive (SRA): PRJNA740342.

## Author Contributions

Design: CAS, TMA, ADJ & RLL. Mass spectrometry experiments and analysis: CAS, MJ & ADJ. NMR experiments and analysis: TMA, ADJ. *In vitro* and *in vivo* biochemical experiments and analysis: CAS. Interpretation of results: CAS, TMA, ADJ & RLL. Wrote the manuscript: CAS & RLL. All authors reviewed and edited the manuscript.

